# Dissecting the effects of GTPase and kinase domain mutations on LRRK2 endosomal localization and activity

**DOI:** 10.1101/2022.10.25.513743

**Authors:** Capria Rinaldi, Christopher S. Waters, Karl Kumbier, Lee Rao, R. Jeremy Nichols, Matthew P. Jacobson, Lani F. Wu, Steven J. Altschuler

## Abstract

Parkinson’s disease-causing LRRK2 mutations lead to varying degrees of Rab GTPase hyperphosphorylation. Puzzlingly, LRRK2 GTPase-inactivating mutations—which do not affect intrinsic kinase activity—lead to higher levels of cellular Rab phosphorylation than kinase-activating mutations. Here, we investigated whether mutation-dependent differences in LRRK2 cellular localization could explain this discrepancy. We discovered that blocking endosomal maturation leads to the rapid formation of mutant LRRK2^+^ endosomes on which LRRK2 phosphorylates substrate Rabs. LRRK2^+^ endosomes are maintained through positive feedback, which mutually reinforces membrane localization of LRRK2 and phosphorylated Rab substrates. Furthermore, across a panel of mutants, cells expressing GTPase-inactivating mutants formed strikingly more LRRK2^+^ endosomes than cells expressing kinase-activating mutants, resulting in higher total cellular levels of phosphorylated Rabs. Our study suggests that an increased probability of LRRK2 GTPase-inactivating mutants to be retained on intracellular membranes over the kinase-activating mutants leads to higher substrate phosphorylation.

## Introduction

Mutations in leucine-rich repeat kinase 2 (LRRK2) are the most prevalent known cause of Parkinson’s disease (PD)^1–3^. LRRK2 comprises several protein domains, including catalytic GTPase and kinase domains, as well as non-catalytic domains believed to mediate protein-protein interactions (Fig. 1A). Disease-causing mutations in the kinase domain, including the most common disease-causing LRRK2 mutation G2019S, can increase the intrinsic (*in vitro*) kinase activity of LRRK2^4–8^. Disease-causing mutations in the GTPase domain, including the highly penetrant LRRK2 mutation R1441G, can decrease the GTPase activity of LRRK2 without altering *in vitro* kinase activity^7–10^. Surprisingly, in cells, these LRRK2 GTPase-inactivating mutations lead to higher phosphorylation levels of its Rab GTPase substrates than the kinase-activating mutations^8,11^.

**Figure 1:**
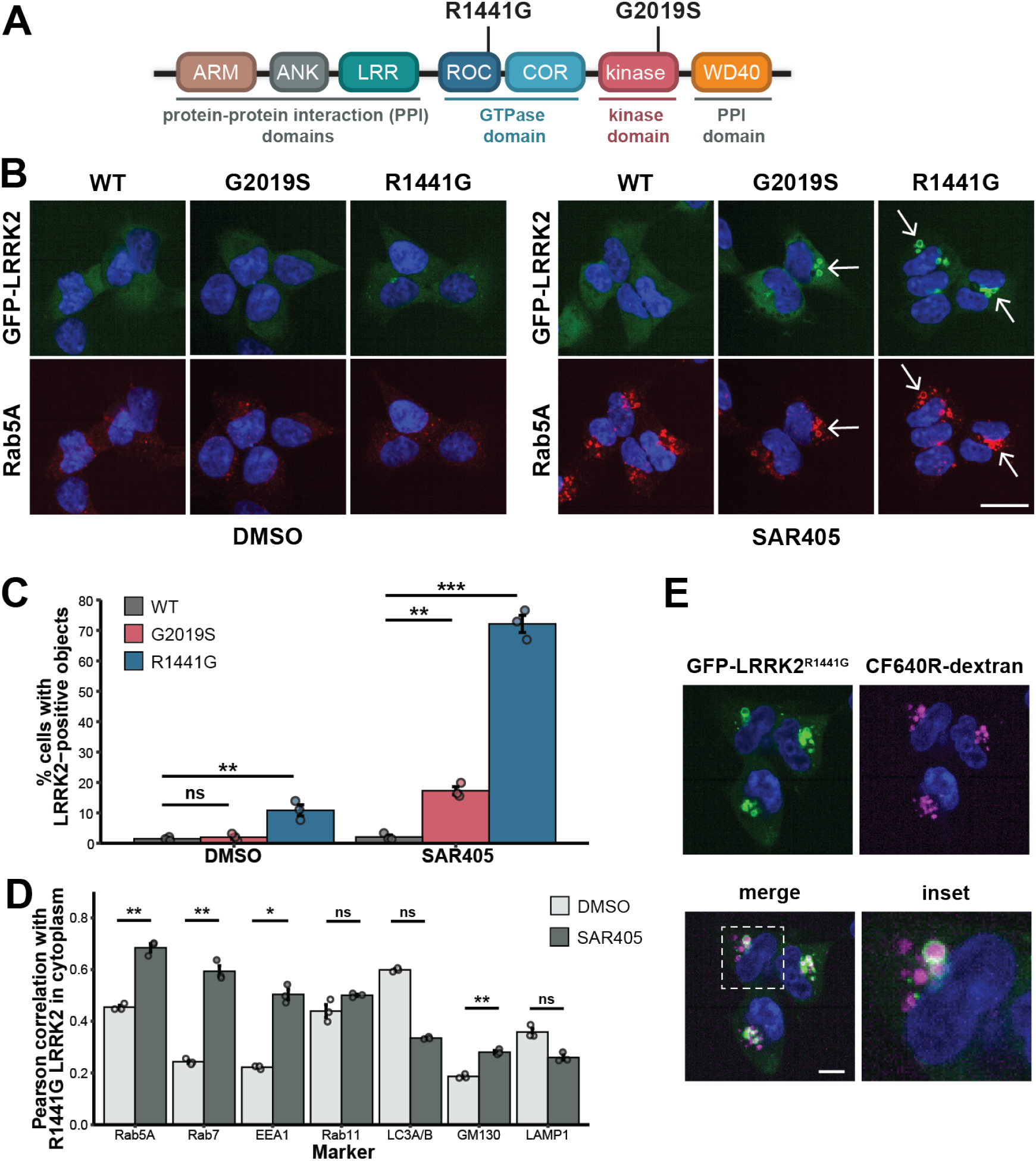
VPS34 inhibition reveals mutant-dependent LRRK2 localization on enlarged endosomes. **(A)** LRRK2 domain structure highlighting two Parkinson’s disease-causing mutations in the Roc-COR GTPase domain (R1441G) and the kinase domain (G2019S). **(B)** Representative images of GFP-LRRK2 variants expressed in stably transfected T-REx HEK293 cells. Cells were treated for 30 min. with DMSO or SAR405 (3 uM) before immunostaining for Rab5A. Arrows indicate localization of LRRK2 mutants on vesicular structures in SAR405. Scale bar, 25 um. **(C)** Quantification of the percent of cells with LRRK2 localized on vesicular structures after 30 min. treatment with DMSO or SAR405 (3 uM). Statistical significance was calculated via one-way ANOVA followed by Dunnett’s post hoc test. (n = 3 wells, >90 cells per well). **(D)** Co-localization analysis of GFP-tagged R1441G LRRK2 with early endosome (Rab5A, EEA1), late endosome (Rab7), recycling endosome (Rab11), autophagosome (LC3A/B), Golgi complex (GM130), or lysosome (LAMP1) markers after 30 min. treatment with DMSO or SAR405 (3 uM). Each point represents the mean of single-cell measurements for all imaged cells within a well. Statistical significance of increased correlation was calculated via unpaired, one-tailed t-test with Bonferroni correction (n = 3 wells, >90 cells per well). **(E)** Representative images of T-REx HEK293 cells expressing GFP-tagged R1441G LRRK2 labeled with 10 kDa CF640R-dextran in the presence of SAR405 (3 uM). Live-cell images shown are from 30 min. after washout of excess dextran to assess localization. Scale bar, 10 um. Error bars: mean +/− SEM. * indicates p values < 0.05; ** indicates p values < 0.01; *** indicates p values < 0.001.

In fact, all disease-causing LRRK2 mutations lead to increased phosphorylation of its Rab GTPase substrates in cells^11,12^. Both LRRK2 and Rab GTPases function on vesicles within the endo-lysosomal system and cycle between cytoplasmic and membrane-bound states^13^. Though largely cytoplasmic, LRRK2 can associate with vesicular membranes to phosphorylate membrane-bound Rabs^14–17^. In some conditions, such as prolonged lysosomal damage, LRRK2 has been shown to be retained on intracellular membranes to a level that can be readily observed by microscopy^16,18–21^. How different LRRK2 mutants affect its membrane association and Rab phosphorylation remains poorly characterized.

Here, we developed a cellular system to dissect the impact of LRRK2 mutations on LRRK2 cellular localization and substrate phosphorylation. We found that acutely blocking endosomal maturation led to the formation of LRRK2-positive endosomes (LRRK2^+^ endosomes) in cells expressing mutant forms of LRRK2. We demonstrated that positive feedback is operating on the LRRK2^+^ endosomes to mutually reinforce the membrane localization of LRRK2 and phosphorylated Rab substrates. Interestingly, across a panel of disease-associated LRRK2 mutants, cell populations expressing GTPase-inactivating mutants formed more LRRK2^+^ endosomes in a dramatically larger fraction of cells than cell populations expressing kinase-activating mutants. Furthermore, the amount of LRRK2^+^ endosomes observed was highly correlated with cellular Rab phosphorylation levels across the panel of mutants. Together, we propose a model wherein LRRK2 GTPase-inactivating mutants have a higher propensity than kinase-activating mutants to form LRRK2^+^ endosomes, thereby providing an explanation for stronger increases in overall cellular Rab phosphorylation.

## Results

### Establishment of a cellular system to monitor, quantify, and compare LRRK2 localization across mutants

To monitor cellular LRRK2 localization, we used a doxycycline-inducible system allowing GFP-tagged LRRK2 plasmids to be stably expressed at relatively low, consistent levels^7,22–24^. We started with three cell lines, expressing WT, G2019S, or R1441G LRRK2 (Fig. 1A). As previously reported, in control conditions LRRK2 expression was largely diffuse and similar between WT and mutant cell lines, though a fraction of R1441G LRRK2 cells contained small cytoplasmic LRRK2 puncta (Fig. 1B)^7,25^.

We searched for a chemical perturbation that would amplify differences in LRRK2 localization across mutants and WT. We reasoned that blocking vesicle trafficking could enrich for a pool of vesicles on which LRRK2 normally signals, thereby amplifying LRRK2 vesicle association events. To test this hypothesis, we evaluated six compounds inhibiting several endo-lysosomal pathways as LRRK2 has functions at multiple stages of vesicle trafficking (Fig. S1A)^26,27^.

Quantification of single-cell phenotypes showed that both chloroquine and SAR405 significantly altered LRRK2 localization for both mutants (Fig. S1A). We chose to further investigate the VPS34 inhibitor SAR405, as this perturbation significantly altered only mutant but not WT LRRK2 localization (Fig. S1A). Upon VPS34 inhibition, the LRRK2 mutant cells had enlarged, LRRK2-positive vesicular structures (Fig. 1B). Strikingly, R1441G compared to G2019S LRRK2-expressing cell populations had a higher proportion of cells containing LRRK2-positive vesicles (72% vs. 17%) (Fig. 1C). Overall, VPS34 inhibition revealed strong differences in enrichment on vesicles between WT and mutant LRRK2, as well as between LRRK2 mutants.

How does VPS34 inhibition affect vesicle trafficking? VPS34 is a phosphoinositide 3-kinase (PI3K) that generates PI(3)P, which promotes maturation of early endosomes^28^. We verified in our system that VPS34 inhibition reduced PI(3)P fluorescence on endosomes (Fig. S1B-C)^29,30^. Furthermore, across WT and mutant LRRK2 cells, VPS34 inhibition increased the total area of objects positive for early endosomal protein Rab5A (Fig. 1B, Fig. S1D)^28,31^. Thus, VPS34 inhibition increases the cellular pool of maturing, Rab5-positive endosomal membranes.

To characterize the LRRK2-positive vesicles induced by VPS34 inhibition, we focused on the strong R1441G phenotype (conclusions are similar for G2019S, data not shown). We found strong co-localization of R1441G LRRK2 with early to late endosome markers (EEA1, Rab5A, Rab7). These strongly co-localized markers had both a high Pearson correlation in VPS34 inhibition conditions and a significant increase in Pearson correlation from DMSO upon VPS34 inhibition (Fig. 1D, Fig. S2A-B). Further, labeled dextran added to the extracellular medium of SAR405-treated cells was present within LRRK2-positive vesicles (Fig. 1E). Overall, we reasoned that the LRRK2-positive vesicles revealed by SAR405 treatment are a subset of enlarged, intermediate vesicles maturing from early (EEA1, Rab5) to late (Rab7) endosomal identity that are capable of transporting endocytic cargo.

Taken together, blocking endosomal maturation via VPS34 inhibition increased the pool of enlarged, maturing endosomes, which revealed differentially amplified LRRK2 endosomal localization across mutants. This provided a setting in which to compare the formation of LRRK2-positive endosomes (LRRK2^+^ endosomes) and downstream consequences across mutants.

### Cells expressing R1441G LRRK2 form more LRRK2^+^ endosomes than cells expressing G2019S LRRK2

We next investigated mutant specific LRRK2^+^ endosome formation in response to acute VPS34 inhibition using video microscopy. For R1441G LRRK2, most cells developed LRRK2^+^ endosomes within 10 minutes. In contrast, for G2019S LRRK2, only a modest proportion of cells with LRRK2^+^ endosomes were observed. WT LRRK2^+^ endosomes were only observed in a very small proportion (1-3%) of cells (Fig. 2A-B, Supp. Video 1-3). The fraction of cells with LRRK2^+^ endosomes was stable over the remainder of the one-hour observation period. Overall, acute VPS34 inhibition induced more rapid LRRK2^+^ endosome formation in a larger proportion of cells expressing R1441G LRRK2 compared to G2019S LRRK2.

**Figure 2:**
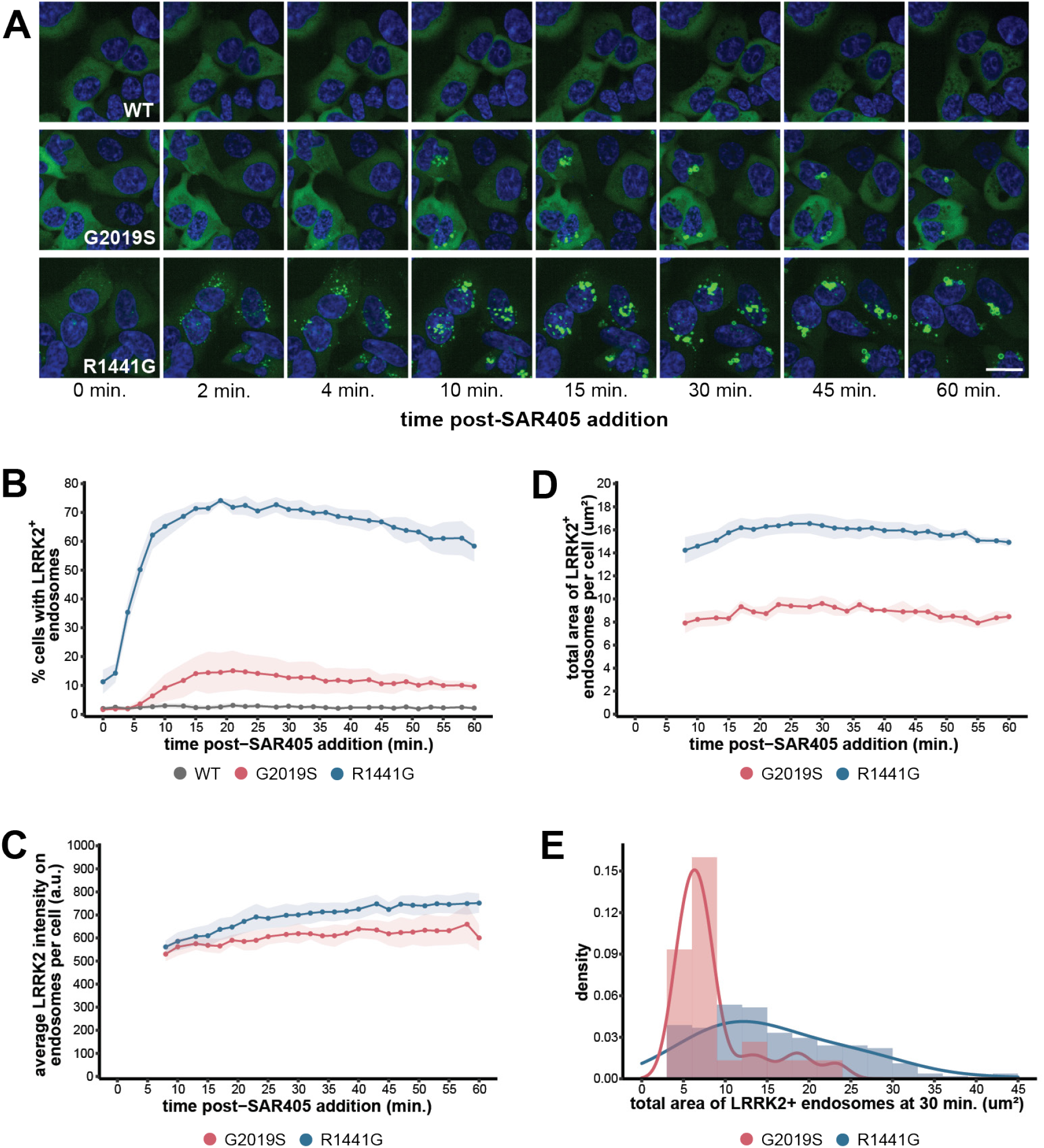
Distinct LRRK2 mutations lead to differential levels of LRRK2^+^ endosome formation. **(A)** Representative sequential images of live T-REx HEK293 cells expressing the indicated GFP-LRRK2 variants treated with SAR405 (3 uM). Cells were imaged approximately every 2 min. Scale bar, 25 um. **(B)** Quantification of percent of cells with LRRK2^+^ endosomes. **(C-D)** Quantification of the average GFP-LRRK2 intensity on endosomes (C) or the total area of LRRK2^+^ endosomes (D) per cell for cells in B containing LRRK2^+^ endosomes. Timepoints in which G2019S LRRK2 was not yet localized to endosomes were omitted. (B-D: Shaded areas: mean +/− SEM, n = 4 independent experiments, >200 cells per LRRK2 genotype in each experiment). **(E)** Single-cell distributions of the total area of LRRK2^+^ endosomes per cell after 30 min. of SAR405 treatment (3 uM). (n = 25 cells G2019S, 181 cells R1441G). (C-E: The number of cells with WT LRRK2 endosome localization was too few for quantification.)

We further examined the subpopulation of cells containing mutant LRRK2^+^ endosomes. After acute VPS34 inhibition, LRRK2^+^ endosomes are first visible as small, punctate structures that become enlarged over time, often resulting from apparent fusion of multiple endosomes (Supp. Video 1-2). The average LRRK2 intensity on endosomes for R1441G LRRK2 was slightly increased compared to G2019S LRRK2 (Fig. 2C) (though quantification at later time points may have been biased due to increased R1441G LRRK2^+^ endosome clustering; Fig. 2A, Supp. Video 1-2). By quantifying the total area of LRRK2^+^ endosomes per cell, we found that R1441G LRRK2 associated with a larger endosomal mass than G2019S LRRK2 (Fig. 2A, Fig. 2D-E).

As controls, we tested that: DMSO treatment did not alter LRRK2 localization (Fig. S1E); SAR405 treatment did not induce significant changes in localization of GFP in cells expressing an empty vector control (Supp. Video 4); and a structurally distinct VPS34 inhibitor, VPS34-IN1, induced similar phenotypes across LRRK2 mutant cell lines (Fig. S1F). In summary, upon acute VPS34 inhibition, the GTPase-inactivating mutant R1441G has a larger area of LRRK2^+^ endosomes in a larger proportion of cells than the kinase-activating mutant G2019S.

### LRRK2^+^ endosome formation leads to amplified cellular Rab phosphorylation levels

Does LRRK2 endosomal membrane localization affect substrate Rab phosphorylation^14,15^? While we observed co-localization of LRRK2 with Rab5A and Rab7 (Fig. 1D), these are not considered validated, endogenous LRRK2 substrates^12^. Thus, we investigated the localization and phosphorylation of two validated LRRK2 phosphorylation substrates, Rab10 and Rab8A^12^.

We first observed the levels and localization of phosphorylated Rab10 (p-Rab10) in cells using IF. Consistent with previous reports, under control conditions, the cellular p-Rab10 signal was increased for cells expressing R1441G LRRK2 and to a lesser extent G2019S LRRK2 (Fig. 3A-B)^11^. After acute VPS34 inhibition, cellular p-Rab10 levels were consistently amplified across the mutant cell populations by ~3-fold, while total Rab10 signal remained largely unchanged (Fig. 3A-B, Fig. S3A-B). Rab10 and p-Rab10 strongly colocalized with R1441G (Fig. 3C, Fig. S3C) and G2019S LRRK2 (data not shown). This colocalization was apparent on LRRK2^+^ endosomes (Fig. 3A, Fig. S3C). Importantly, the levels of p-Rab10 decreased to baseline after treatment with the LRRK2 kinase inhibitor MLi-2, confirming that the observed increases in phosphorylation were due to LRRK2 kinase activity^12^ (Fig. 3A-B). Similar mutant-specific trends and MLi-2 effects were observed for the substrate Rab8A (using Western blot to measure p-Rab8A levels and IF to observe localization of total Rab8A; Fig. 3C-D, Fig. S3C).

**Figure 3:**
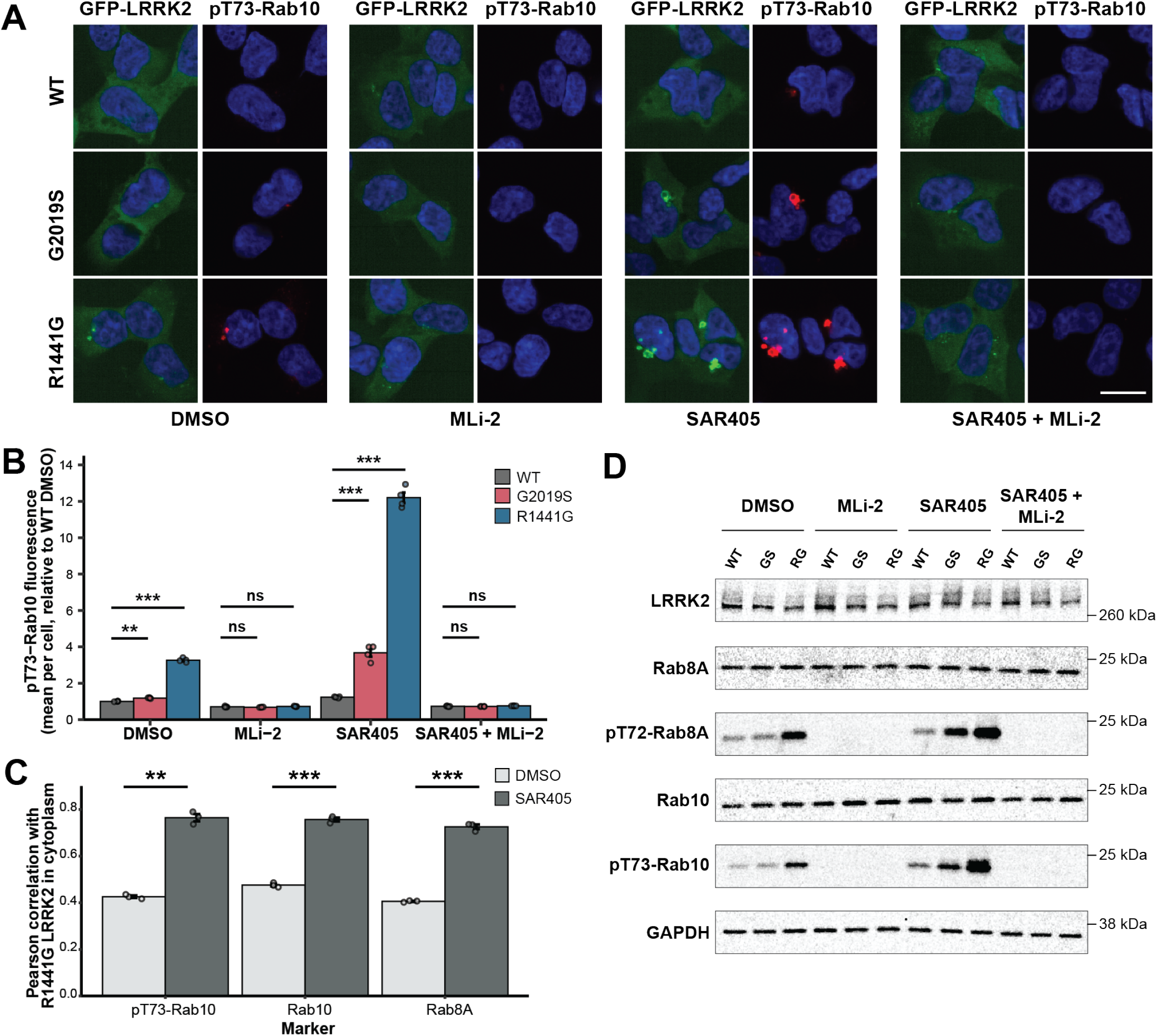
LRRK2 phosphorylates Rab10 on LRRK2^+^ endosomes. **(A)** Representative images of T-REx HEK293 cells expressing GFP-LRRK2 variants after 30 min. drug treatment and immunostaining for LRRK2-specific phosphosite on pRab10 (pT73-Rab10). MLi-2: 1 uM, SAR405: 3 uM. Scale bar, 20 um. **(B)** Quantification of mean pT73-Rab10 fluorescence intensity per cell displayed as fold change relative to DMSO-treated cells expressing WT LRRK2. Each point represents the mean of single-cell measurements for all imaged cells within a well. Statistical significance was calculated via one-way ANOVA followed by Dunnett’s post hoc test (n = 4 wells, >200 cells per well). **(C)** Co-localization analysis of GFP-tagged R1441G LRRK2 with pT73-Rab10, Rab10, or Rab8A after 30 min. treatment with DMSO or SAR405 (3 uM). Each point represents the mean of single-cell measurements for all imaged cells within a well. Statistical significance was calculated via unpaired, one-tailed t-test with Bonferroni correction (n = 3 wells, >130 cells per well). **(D)** Western blot for phosphorylated Rab8A and Rab10 after 30 min. drug treatment. MLi-2: 1 uM, SAR405: 3 uM. Error bars: mean +/− SEM. ** indicates p values < 0.01; *** indicates p values < 0.001.

Furthermore, in cells with mutant LRRK2^+^ endosomes after VPS34 inhibition, cells with higher cellular p-Rab10 levels largely had a greater LRRK2^+^ endosomal mass (Fig. S3D). Thus, the differences in p-Rab10 levels across cell populations reflect both the proportion of cells with LRRK2^+^ endosomes and the mass of LRRK2^+^ endosomes within single cells on which high levels of p-Rab10 are observed. Overall, VPS34 inhibition revealed amplification of LRRK2 and phosphorylated substrate levels on the same LRRK2^+^ endosomal membranes.

### LRRK2^+^ endosomes are established and maintained through Rab phosphorylation

We next wondered whether Rab phosphorylation could play a role in maintaining LRRK2 on the membrane of LRRK2^+^ endosomes, as strong binding of LRRK2 to phosphorylated Rabs was recently described *in vitro* on lipid bilayers^32^.

First, we investigated the role of Rab phosphorylation in the establishment of mutant LRRK2^+^ endosomes. Co-treatment of cells for 30 minutes with both the VPS34 inhibitor and the LRRK2 kinase inhibitor MLi-2 blocked the formation of LRRK2^+^ endosomes (Fig. S4A). To further validate this result, we overexpressed the phosphatase PPM1H, which dephosphorylates Rab proteins and counteracts LRRK2 signaling^33^. Overexpression of PPM1H dramatically decreased the proportion of cells with LRRK2^+^ endosomes formed upon VPS34 inhibition for both R1441G and G2019S LRRK2 (Fig. 4A-B, Fig. S4B). (We confirmed that PPM1H overexpression decreased p-Rab10 levels without affecting total Rab10 levels and that overexpression of the inactive H153D PPM1H mutant had no effect on R1441G LRRK2 localization; Fig. 4B, Fig. S4C.) Thus, Rab dephosphorylation inhibits the establishment of LRRK2^+^ endosomes.

**Figure 4:**
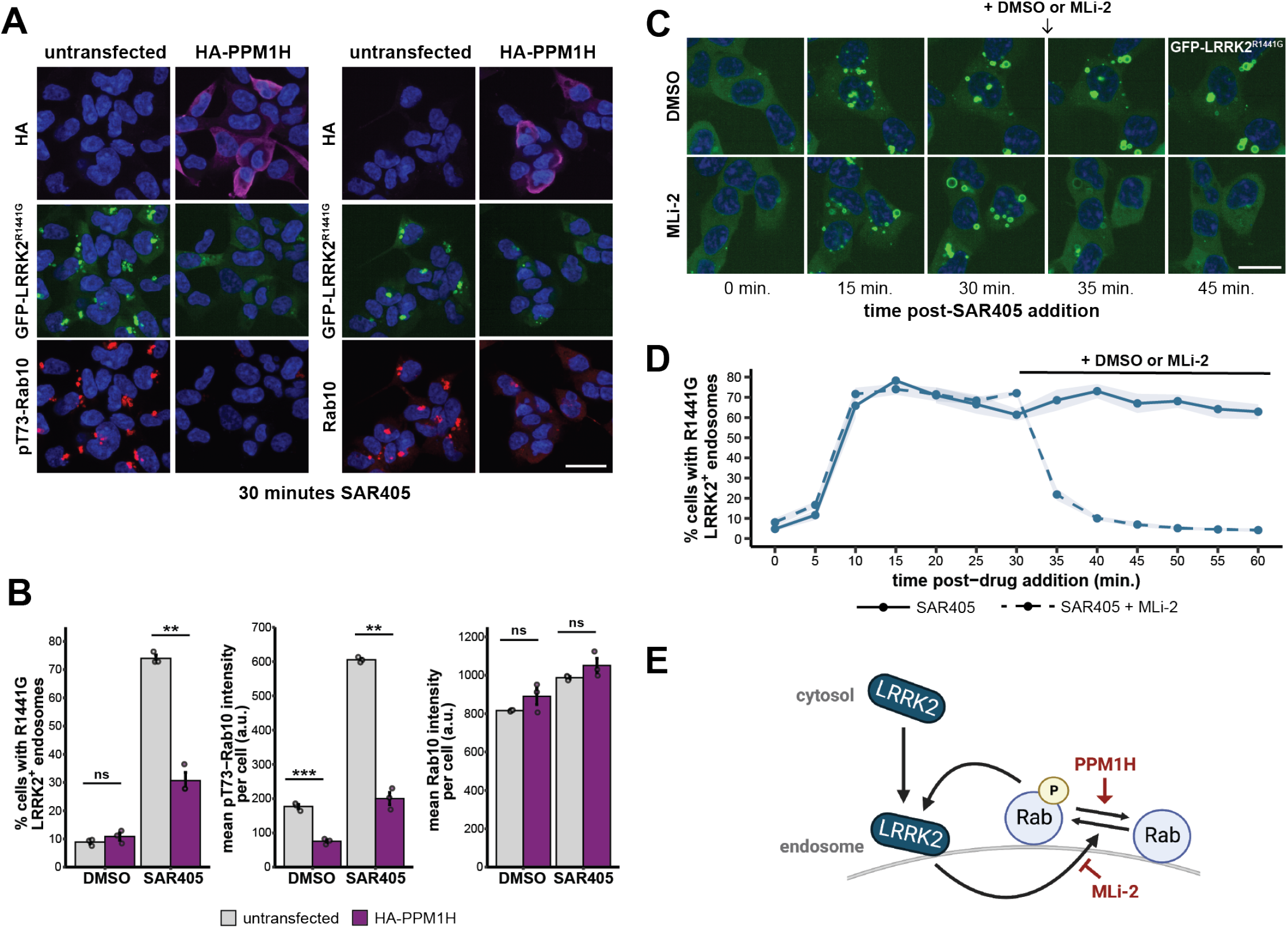
LRRK2^+^ endosomes are established and maintained through Rab phosphorylation. **(A)** Representative images of stable T-REx HEK293 cells expressing GFP-tagged R1441G LRRK2 and transiently transfected with a plasmid encoding HA-PPM1H. Cells were treated for 30 min. with SAR405 (3 uM) before immunostaining for HA-tag and total Rab10 or pT73-Rab10. Scale bar, 30 um. **(B)** Left: percent of cells with R1441G LRRK2^+^ endosomes. Middle and right: mean pT73-Rab10 or mean total Rab10 fluorescence intensity per cell. Statistical significance was calculated via unpaired, two-tailed t-test with Bonferroni correction (n = 3 wells, >250 cells per well). Error bars: mean +/− SEM. ** indicates p values < 0.01; *** indicates p values < 0.001. **(C)** Representative sequential live-cell images of stable T-REx HEK293 cells expressing GFP-tagged R1441G LRRK2 treated with SAR405 (3 uM). After 30 min., DMSO or MLi-2 (1 uM) was added to the cell media. Cells were imaged approximately every 5 min. Scale bar, 20 um. **(D)** Quantification of the percent of cells with LRRK2^+^ endosomes. Shaded areas: mean +/− SEM (n = 7 fields of view per condition, >20 cells analyzed per field of view). **(E)** Model of positive feedback on endosomes. Red text: perturbations used in A-D.

Second, we investigated the role of Rab phosphorylation in the maintenance of mutant LRRK2^+^ endosomes. After 30 minutes of VPS34 inhibition to enrich R1441G LRRK2 on endosomes, we added DMSO or MLi-2 to the cell media and monitored LRRK2^+^ endosomes via live imaging (Fig. 4C). After MLi-2 treatment, VPS34 inhibitor-induced LRRK2^+^ endosomes were rapidly lost for both R1441G and G2019S LRRK2 (Fig. 4C-D, Fig. S4D). Thus, the ability of LRRK2 to phosphorylate Rabs is required for the maintenance of mutant LRRK2^+^ endosomes once established. Together, our results suggest that positive feedback is operating on LRRK2^+^ endosomes to maintain LRRK2 and Rab localization (Fig. 4E).

### Differential effects of GTPase-inactivating and kinase-activating mutations on LRRK2 localization and activity are consistent across a panel of mutants

Finally, to test the generality of our findings for G2019S and R1441G LRRK2, we expanded our study to include a total of 15 LRRK2 genotypes (WT LRRK2, 11 disease-associated LRRK2 mutants, protective variant R1398H LRRK2, kinase dead D2017A LRRK2, and GTP binding-deficient T1348N LRRK2) (Fig. 5A). The selected mutations have a range of reported effects on LRRK2 activity, including modifications to the protein’s GTPase and kinase functions (Supp. Table 1). For each mutant, we quantified both the proportion of cells with LRRK2^+^ endosomes and cellular p-Rab10 levels upon acute VPS34 inhibition (Fig. 5B-D).

**Figure 5:**
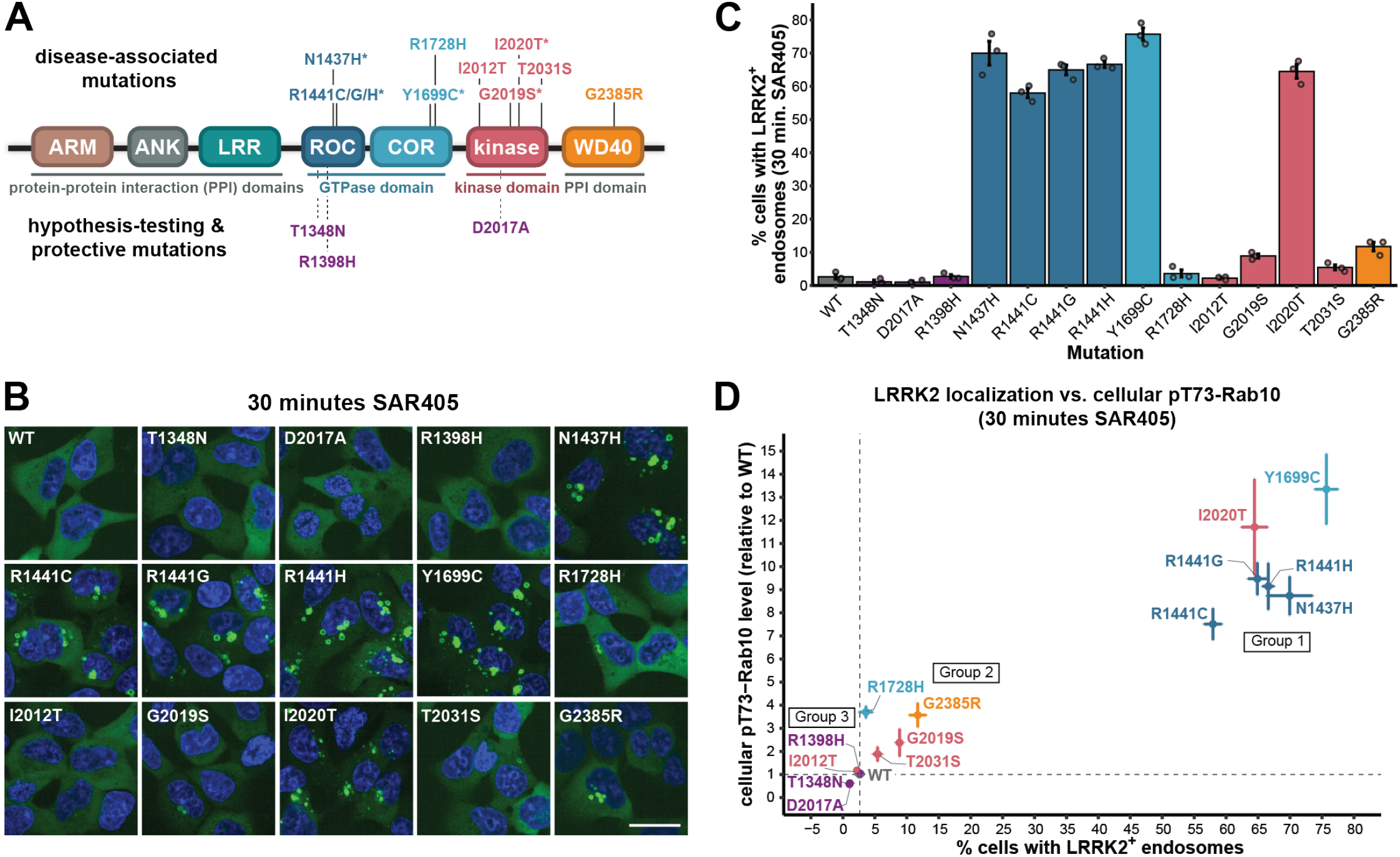
GTPase-inactivating LRRK2 mutants have higher LRRK2^+^ endosome formation and cellular p-Rab10 levels than kinase-activating mutants. **(A)** LRRK2 domain structure denoting the location of all mutations analyzed. Disease-associated mutations are on the top, while hypothesis-testing mutations (T1348N, D2017A) and a protective mutation (R1398H) are on the bottom. Asterisks indicate mutations that are causal for PD. **(B)** Representative live-cell images of T-REx HEK293 cells expressing the indicated GFP-LRRK2 variants after 30 min. treatment with SAR405 (3 uM). Scale bar, 25 um. **(C)** Quantification of the percent of cells with LRRK2^+^ endosomes observed via live imaging after 30 min. SAR405 treatment (3 uM) (n = 3 independent experiments, >500 cells analyzed per LRRK2 genotype in each experiment). **(D)** Scatter plot showing the percent of cells with LRRK2^+^ endosomes (using data from C) vs. cellular pT73-Rab10 levels after 30 min. SAR405 treatment (3 uM). pT73-Rab10 levels were quantified as the mean pT73-Rab10 fluorescence intensity per cell via immunofluorescence (n = 3 independent experiments, >550 cells analyzed per LRRK2 genotype in each experiment). Dashed lines indicate values for WT LRRK2. Group 1: Y1699C, I2020T, R1441C/G/H, N1437H. Group 2: G2385R, G2019S, T2031S, R1728H. Group 3: R1398H, I2012T, T1348N, D2017A. Error bars: mean +/− SEM.

We note that p-Rab10 levels across LRRK2 mutants in control conditions were consistent with previously reported values (Fig. S5A)^8^, and as before (Fig. 3B), p-Rab10 levels were increased ~3-fold across mutants in VPS34-inhibited vs. untreated conditions (Fig. S5B).

The strong correlation we observed between LRRK2^+^ endosome formation and p-Rab10 levels suggested three groupings of mutants, depending on whether LRRK2^+^ endosome formation and p-Rab10 levels were well above, slightly above or close to WT LRRK2 levels (Groups 1-3, Fig. 5D). Group 1 consists of disease-causing mutants that highly increase the formation of LRRK2^+^ endosomes and lead to highly increased p-Rab10 levels (Y1699C, I2020T, R1441C/G/H, and N1437H). All Group 1 mutants, with the exception of I2020T in the kinase domain, have reduced GTPase activity and increased GTP binding (Supp. Table 1). Furthermore, all Group 1 mutants are dephosphorylated at Ser935, which disrupts LRRK2 interaction with 14-3-3 proteins^7,22^ (Supp. Table 1).

Group 2 contains disease-associated mutants that only moderately increase the formation of LRRK2^+^ endosomes and modestly increased p-Rab10 levels (R1728H, T2031S, G2019S, and G2385R). Overall, mutants in Group 2 have increased *in vitro* kinase activity, though G2385R has conflicting reported activity (Supp. Table 1).

Finally, Group 3 is made up of mutants that have LRRK2^+^ endosome and p-Rab10 levels that are similar, or decreased, compared to WT (I2012T, R1398H, D2017A, and T1348N). I2012T (a putative disease allele with decreased kinase activity) and R1398H (a putative protective allele with increased GTPase activity and decreased GTP binding) behaved similarly to WT. D2017A (kinase dead) and T1348N (no GTP binding) had minimal LRRK2^+^ endosomes and low p-Rab10 levels.

Across the panel of LRRK2 mutants, we found that GTPase-inactivating mutants consistently had stronger effects on LRRK2 membrane localization and Rab phosphorylation compared to kinase-activating LRRK2 mutants. Furthermore, as demonstrated by our results for Group 3 mutants, decreases in LRRK2 GTP binding or kinase activity reduce LRRK2^+^ endosome formation to WT levels or lower. Overall, the connection between LRRK2 membrane localization and Rab phosphorylation provides an explanation for reported inconsistencies in relative p-Rab levels across mutants in cellular and *in vitro* assays, which cannot capture effects of LRRK2 localization (Fig. S5C).

## Discussion

All PD-causing LRRK2 mutations lead to substrate hyperphosphorylation in cells, albeit to varying degrees. Here, we investigated mechanisms underlying how LRRK2 GTPase-inactivating mutations—which do not affect kinase activity *in vitro*—could lead to much higher substrate phosphorylation than kinase-activating mutations in cells. We found that GTPase-inactivating LRRK2 mutants have a higher propensity than kinase-activating mutants to be maintained on Rab-containing endosomal membranes, thereby allowing for greater levels of Rab phosphorylation.

Acute VPS34 inhibition enriches for a pool of enlarged, maturing endosomes by delaying subsequent maturation stages^28,30,34^. This enrichment allowed us to visualize strong recruitment of LRRK2 to endosomes and detect mutant-dependent differences. We speculate that VPS34 inhibition prolongs an endosomal stage at which LRRK2 signaling normally occurs at lower levels. Prior studies described WT LRRK2 signaling on endosomes and, in one case, demonstrated that WT LRRK2 could regulate VPS34-dependent endosomal maturation^17,35–37^. Interestingly, VPS34 depletion causes neuronal degeneration in mice by disrupting the endosomal system, and multiple risk loci for PD are related to phosphoinositide signaling^38–40^. Furthermore, LRRK2 has been reported to aberrantly localize to enlarged endosomal structures in brain tissue from patients with dementia with Lewy bodies^41^, and alterations in early endosomal morphology and endocytic cargo processing have been found in PD patient brain tissue^42^. Future studies may explore whether VPS34 inhibition mirrors disease-relevant cellular states, such as aging-related changes in phosphoinositide pools.

The establishment and maintenance of LRRK2^+^ endosomes in our system is consistent with a model of positive feedback operating on endosomal membranes (Fig. 4E). First, LRRK2 associates with endosomes^17,37^, which occurs with higher probability under conditions of VPS34 inhibition. Second, phosphorylation of Rabs by LRRK2 inhibits Rab membrane disassociation, leading to accumulation of p-Rabs^11,13,14^. Third, LRRK2 binds strongly to p-Rabs, leading to membrane accumulation of LRRK2 through a feed-forward pathway^32^. Together, LRRK2 and Rabs mutually and positively enhance the endosomal membrane localization of one another^32^. Consistent with a model of overall positive feedback, we found that enhancing Rab dephosphorylation (via PPM1H expression) and inhibiting the ability of LRRK2 to phosphorylate Rabs (via MLi-2 treatment) blocked LRRK2^+^ endosome establishment and maintenance. Our study leaves open whether this positive feedback operates on other cellular membranes where LRRK2 can localize (such as at the lysosome or Golgi), as well as where and to what degree this feedback occurs in untreated cells.

This feedback model provides a framework in which to suggest mutant-specific mechanisms underlying differences in cellular p-Rab levels. For example, the kinase-activating mutants may have increased ability to perpetuate feedback through Rab phosphorylation (due to their increased intrinsic kinase activity)^4,8^, while the GTPase-inactivating mutants may have increased probability to initiate feedback through increased initial membrane association (similar to WT LRRK2 in the GTP-bound state or through reduced 14-3-3 binding in the cytoplasm)^4,7,43^. Consistent with this model, LRRK2 GTPase-inactivating mutants are recruited more efficiently to the Golgi by Rab29 than WT LRRK2^16^, and the mutant with the strongest membrane association phenotype (Y1699C LRRK2; Fig. 5D) has both decreased GTPase activity and increased *in vitro* kinase activity^8,44,45^. Ultimately, our data show that GTPase-inactivating mutants have much stronger net effects than kinase-activating mutants on this positive feedback as measured by increases in both LRRK2^+^ endosomal mass and p-Rab10 levels. Thus, even without elevated intrinsic kinase activity, the observed kinase output of GTPase-inactivating mutants in cells can be higher than that of kinase-activating mutants. Further elucidation of this observation will be aided by quantitative parameterization of this feedback circuit in cells, including measured on-, off- and feedback-rates.

LRRK2 possesses additional layers of intramolecular regulation involving both kinase and GTPase activity not studied here. For example, LRRK2 exists both in monomer (primarily cytoplasmic) and dimer/oligomer (enriched at membranes) states^43,46^. The LRRK2 monomer-dimer cycle is thought to be regulated via GTP binding, and LRRK2 phosphorylates Rabs only in the active, dimeric/oligomeric state^4,43,47,48^. Furthermore, LRRK2 contains numerous autophosphorylation sites across domains with incompletely understood functions^4,49,50^. Future studies may explore the role of autophosphorylation sites within the GTPase domain of LRRK2 in regulating LRRK2^+^ endosome formation.

We observed a qualitative but striking correlation of disease risk with the three LRRK2 mutant phenotypic groupings (defined by high, medium and low levels of LRRK2^+^ endosomes and Rab phosphorylation; Fig. 5D). All mutations in Group 1 (largely GTPase-inactivating mutations) are causal for PD, with estimates of disease penetrance at age 80 being high for R1441C/G mutation carriers (>80%). In Group 2 (largely kinase-activating mutations), G2019S is considered causative, albeit with a lower penetrance (25% - 42.5%), and the remaining mutations modestly increase risk of disease. Finally, R1398H in Group 3, which increases GTPase activity, is a protective variant for PD^1,51–53^. Thus, LRRK2 cellular localization, which is functionally connected to cellular Rab phosphorylation, may be a driver of downstream disease risk.

Our study provides a rationale for why disease-associated mutations across LRRK2 lead to differing degrees of substrate phosphorylation in cells. In cells expressing GTPase-inactivating mutants, both LRRK2 endosomal membrane localization and downstream Rab phosphorylation are strongly increased compared to kinase-activating mutants. It is encouraging that Rab dephosphorylation and LRRK2 kinase inhibition reversed aberrant localization of LRRK2 mutants on endosomal membranes, as LRRK2 kinase inhibition is currently being evaluated in clinical trials for PD^54^. Our results also suggest that strategies to promote the cytoplasmic localization of LRRK2, perhaps by modulating GTPase function or disrupting LRRK2 dimerization, could be valuable alternative methods to explore for LRRK2 PD treatment.

---

**Figure S1:**
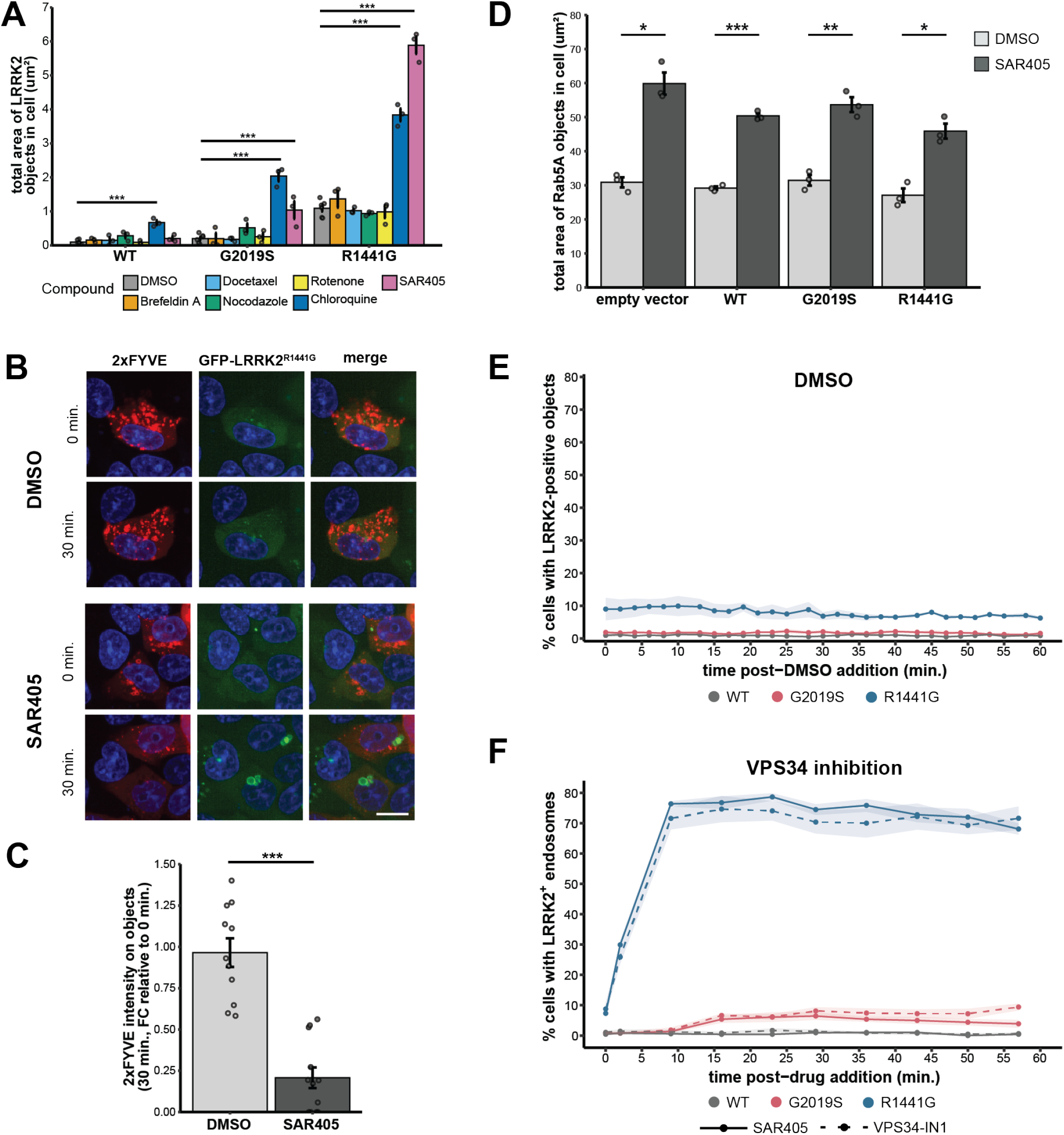
VPS34 inhibition reveals mutant-specific LRRK2 localization pattern. **(A)** Quantification of the mean total area of LRRK2 localized on objects per cell. Each point represents the mean of single-cell measurements for all imaged cells within a well. Statistical significance was calculated via one-way ANOVA followed by Dunnett’s post hoc test (n = 6 wells for DMSO, 3 wells for all other compounds, >65 cells per well). **(B)** Representative images of live T-REx HEK293 cells expressing GFP-tagged R1441G LRRK2 and transiently transfected with a plasmid encoding the PI3P probe mCherry-2xFYVE. Cells were treated with DMSO or SAR405 (3 uM) before live imaging. SAR405 treatment reduced mCherry-2xFYVE signal concomitant with an increase in R1441G LRRK2 endosomal membrane localization. Scale bar, 15 um. **(C)** Quantification of mCherry-2xFYVE intensity on objects as a percent of total cellular mCherry-2xFYVE intensity in cells expressing GFP-R1441G LRRK2. Data shown as fold change (FC) in 2xFYVE intensity on objects 30 min. after compound treatment relative to 2xFYVE intensity on objects in the same cell prior to treatment. Statistical significance was calculated via unpaired, two-tailed t-test (n = 11 cells DMSO, 12 cells SAR405). **(D)** Quantification of total area of Rab5A-positive objects after 30 min. treatment with DMSO or SAR405 (3 uM). Each point represents the mean of single-cell measurements for all imaged cells within a well. Statistical significance was calculated via unpaired, two-tailed t-tests with Bonferroni correction (n = 3 wells, >90 cells per well). A-D: Error bars: mean +/− SEM. * indicates p values <0.05; ** indicates p values < 0.01; *** indicates p values < 0.001. **(E)** T-REx HEK293 cells expressing the indicated GFP-tagged LRRK2 variants were treated with DMSO and analyzed via live-cell imaging. Cells were imaged approximately every 2 min. Quantification of percent of cells with LRRK2 localized on objects. (n = 4 independent experiments, >200 cells analyzed per LRRK2 genotype in each experiment). **(F)** T-REx HEK293 cells expressing the indicated GFP-tagged LRRK2 variants were treated with SAR405 (3 uM) or VPS34-IN1 (3 uM) and analyzed via live-cell imaging. Cells were imaged approximately every 7 min. Quantification of percent of cells with LRRK2^+^ endosomes. (n = 8 fields of view per condition, >20 cells analyzed per field of view). E-F: Shaded areas: mean +/− SEM.

**Figure S2:**
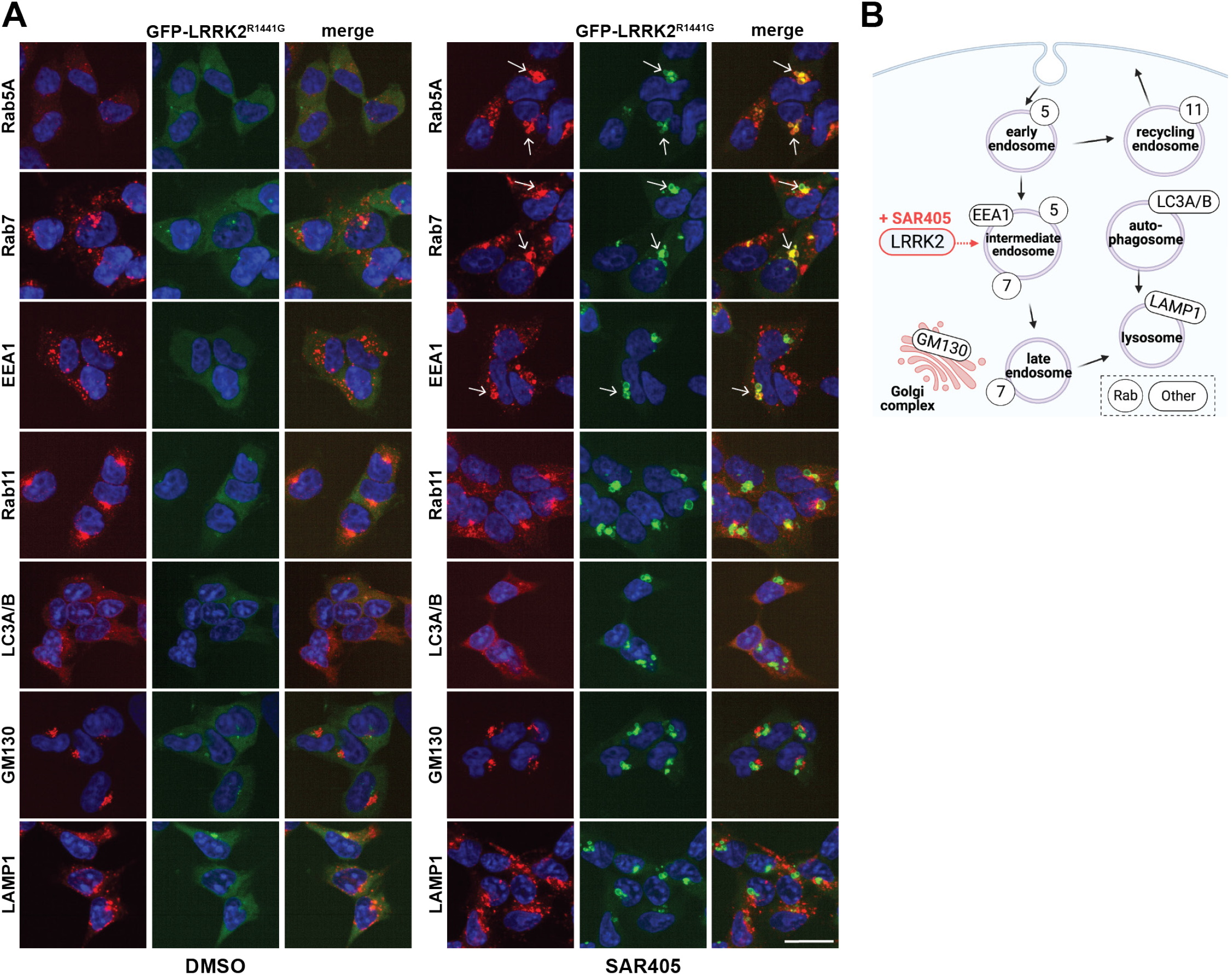
LRRK2 co-localizes with early and late endosomal markers upon VPS34 inhibition. **(A)** Representative images demonstrating co-localization of GFP-tagged R1441G LRRK2 with endo-lysosomal biomarkers. Cells were treated for 30 min. with DMSO or SAR405 (3 uM) before immunostaining for indicated proteins. Arrows indicate strong co-localization of R1441G LRRK2-positive vesicles with Rab5A, Rab7, and EEA1 in the SAR405 condition. Scale bar, 25 um. **(B)** Simplified cartoon indicating the identity of vesicles or organelles labeled with the endo-lysosomal biomarkers^55–57^. SAR405 treatment reveals LRRK2 co-localization with markers of intermediate, maturing endosomes.

**Figure S3:**
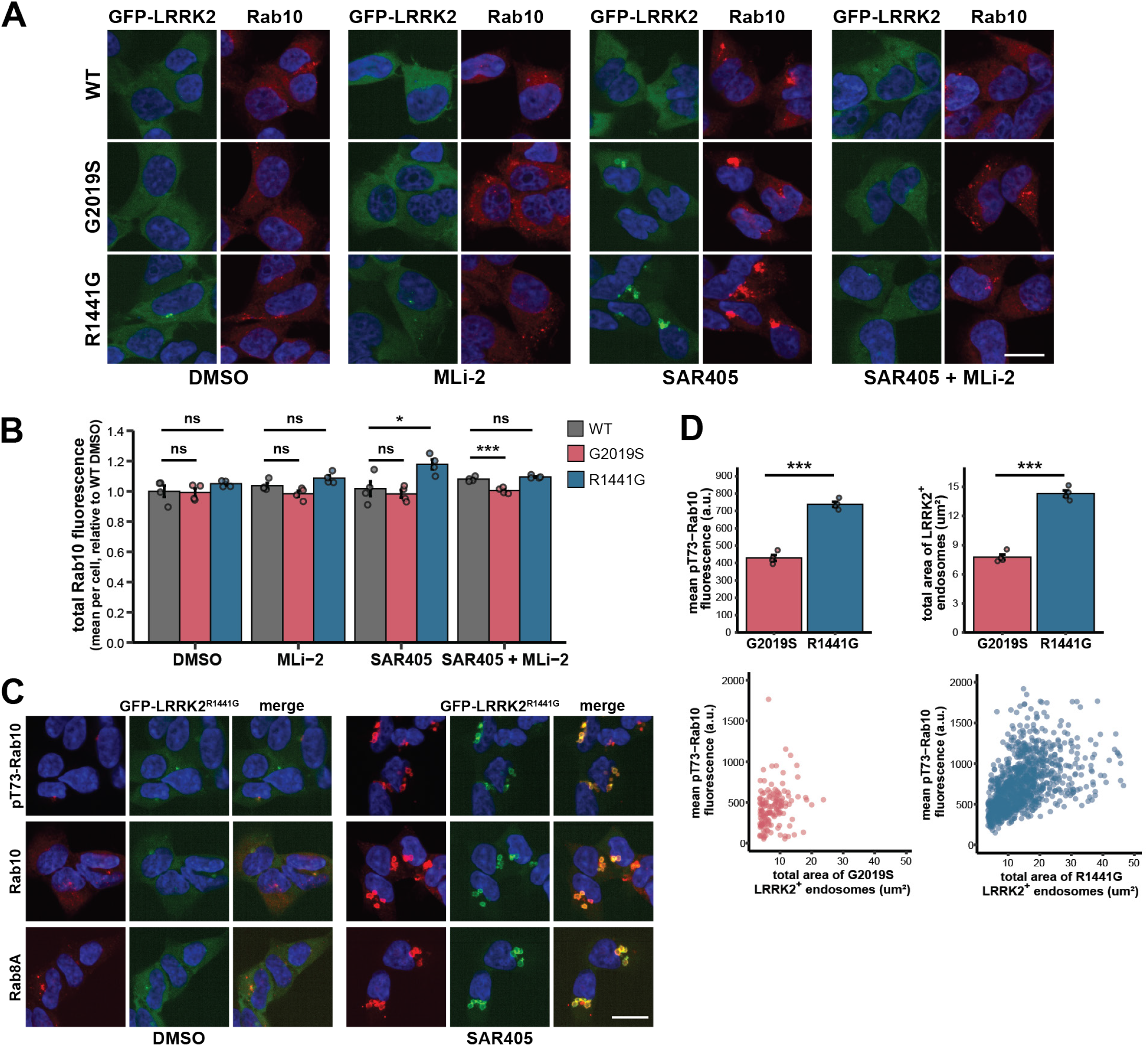
Effects of VPS34 inhibition on total Rab10 intensity, pT73-Rab10 intensity and LRRK2^+^ endosomes in single cells, and LRRK2 co-localization with Rab substrates. **(A)** Representative images of T-REx HEK293 cells expressing GFP-tagged LRRK2 variants after 30 min. drug treatment and immunostaining for total Rab10. MLi-2: 1 uM, SAR405: 3 uM. Scale bar, 20 um. **(B)** Quantification of mean total Rab10 fluorescence intensity per cell displayed as fold change relative to DMSO-treated cells expressing WT LRRK2. Each point represents the mean of single-cell measurements for all imaged cells within a well. Statistical significance was calculated via one-way ANOVA followed by Dunnett’s post hoc test (n = 4 wells, >250 cells per well). * indicates p values < 0.05; *** indicates p values < 0.001. **(C)** Representative images demonstrating strong co-localization of GFP-tagged R1441G LRRK2 with total and phospho-Rab proteins upon VPS34 inhibition. Cells were treated for 30 min. with DMSO or SAR405 (3 uM) before immunostaining for Rab10, pT73-Rab10, or Rab8A. Scale bar, 20 um. **(D)** Top: Quantification of the mean pT73-Rab10 fluorescence or total area of LRRK2^+^ endosomes per cell for SAR405-treated cells in Fig. 3B containing LRRK2^+^ endosomes. Statistical significance was calculated via unpaired, two-tailed t-tests with Bonferroni correction (n = 4 wells, >25 cells G2019S and >220 cells R1441G analyzed per well). Bottom: Scatter plots of mean pT73-Rab10 fluorescence vs. total area of LRRK2^+^ endosomes per cell for pooled single cells summarized in above barplots. (n = 144 cells G2019S, 1051 cells R1441G). Error bars: mean +/− SEM.

**Figure S4:**
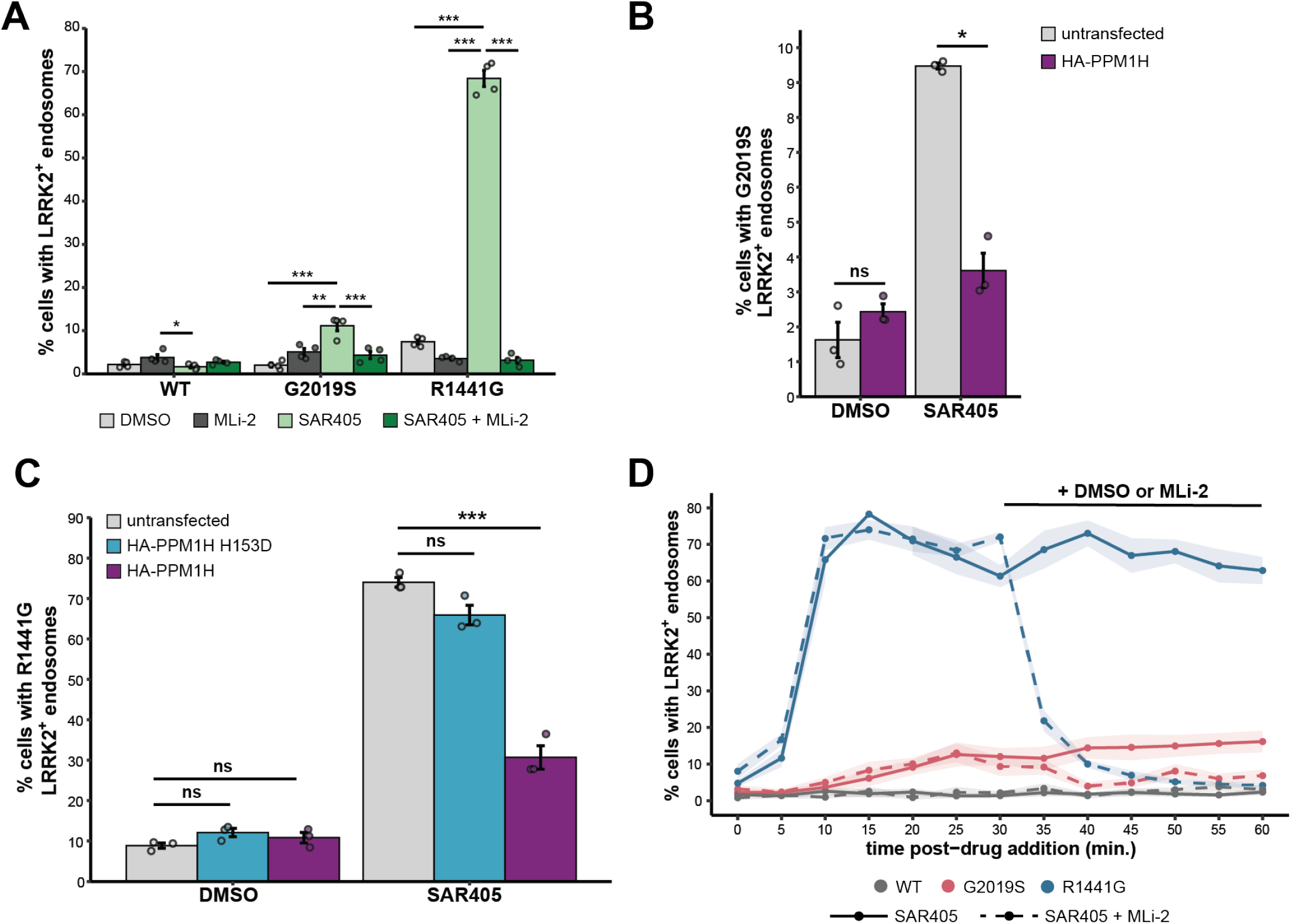
Additional analysis of PPM1H and PPM1H H153D overexpression and LRRK2 kinase inhibition effects on LRRK2^+^ endosome formation. **(A)** T-REx HEK293 cells expressing GFP-LRRK2 variants were treated for 30 min. with the indicated compounds before fixation and quantification of the percent of cells with LRRK2^+^ endosomes. MLi-2: 1 uM, SAR405: 3 uM. Statistical significance was calculated via one-way ANOVA followed by Tukey’s post hoc test (n = 4 wells, >200 cells per well). * indicates p values < 0.05; ** indicates p values <0.01; *** indicates p values < 0.001, unlabeled comparisons were not significant. **(B)** Stable T-REx HEK293 cells expressing GFP-tagged G2019S LRRK2 were transiently transfected with a plasmid encoding HA-PPM1H. Cells were treated for 30 min. with DMSO or SAR405 (3 uM) before immunostaining for HA tag and quantifying the percent of untransfected or PPM1H-positive cells with LRRK2^+^ endosomes. Statistical significance was calculated via unpaired, two-tailed t-test with Bonferroni correction (n = 3 wells, >450 cells per well). **(C)** T-REx HEK293 cells expressing GFP-tagged R1441G LRRK2 were transiently transfected with a plasmid encoding either HA-PPM1H or the inactive mutant HA-PPM1H H153D. Cells were treated for 30 min. with DMSO or SAR405 (3 uM) before immunostaining for HA tag and quantifying the percent of untransfected or PPM1H-positive cells with LRRK2^+^ endosomes. Statistical significance was calculated via one-way ANOVA followed by Dunnett’s post hoc test (n = 3 wells, >300 cells per well). A-B: Error bars: mean +/− SEM. * indicates p values < 0.05; *** indicates p values < 0.001. **(D)** Quantification of LRRK2^+^ endosomes in T-REx HEK293 cells expressing GFP-tagged LRRK2 variants treated with SAR405 (3 uM) and monitored via live imaging. After 30 min., DMSO or MLi-2 (1 uM) was added to the cell media. Cells were imaged approximately every 5 min. Data for R1441G is the same as in Fig. 5D. Shaded areas: mean +/− SEM (n = 7 fields of view per condition, >20 cells analyzed per field of view).

**Figure S5:**
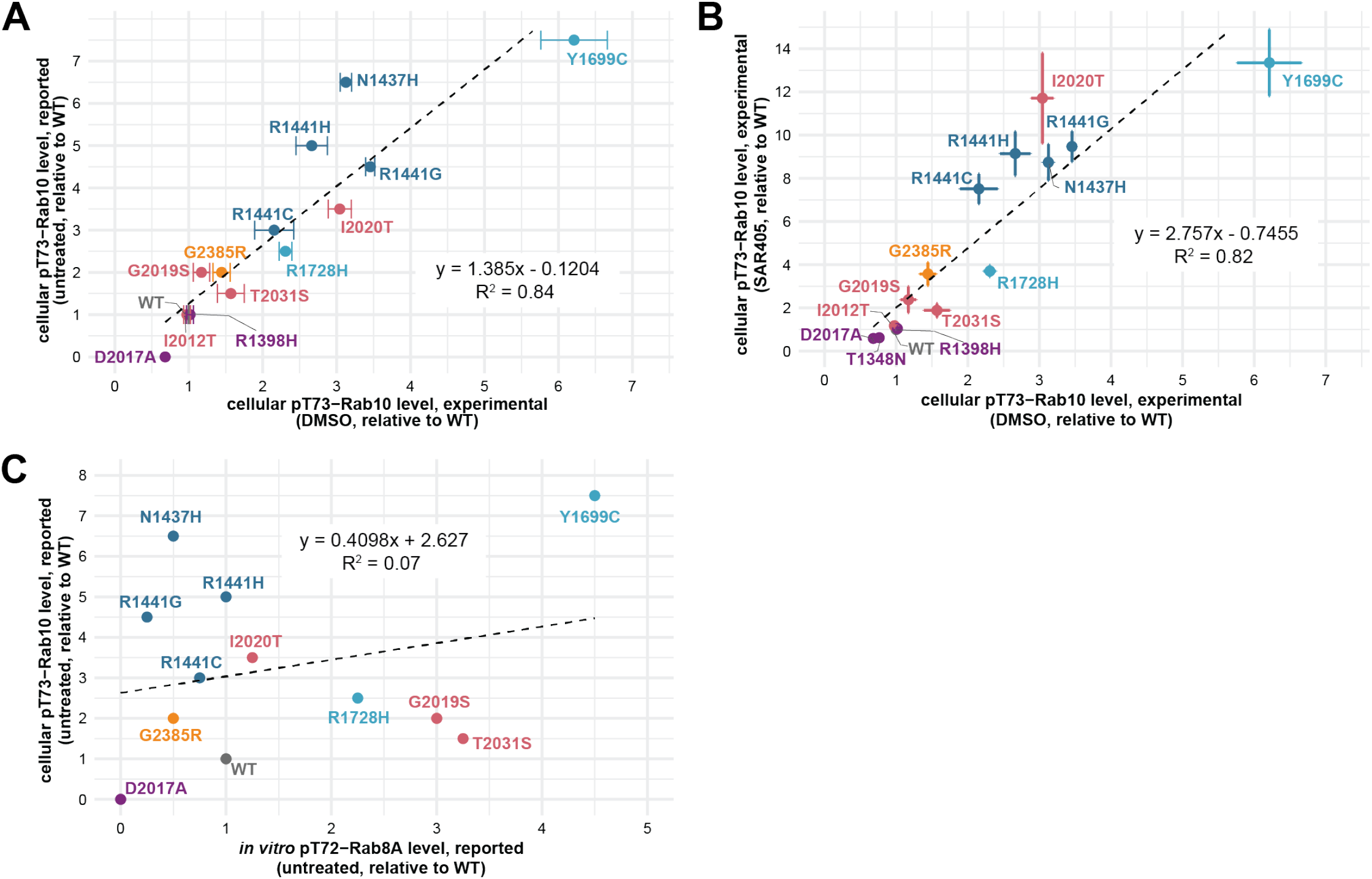
Relationships between experimental data and reported data for *in vitro* and cellular Rab phosphorylation. **(A)** Scatter plot demonstrating that cellular p-Rab10 levels measured in this work in control DMSO conditions correlate with reported cellular p-Rab10 levels in untreated conditions. Cellular p-Rab10 levels on x-axis were quantified as the mean pT73-Rab10 fluorescence intensity per cell via immunofluorescence after 30 min. treatment with DMSO (n = 3 independent experiments, >600 cells analyzed per LRRK2 genotype in each experiment). Error bars: mean +/− SEM. Cellular p-Rab10 levels on y-axis were estimated from Figure 1 of Kalogeropulou et al^8^. **(B)** Scatter plot demonstrating that SAR405 treatment amplifies cellular p-Rab10 levels approximately 3-fold (slope = 2.8) across the mutations analyzed. Cellular p-Rab10 levels were quantified as the mean pT73-Rab10 fluorescence intensity per cell via immunofluorescence after 30 min. treatment with either DMSO or SAR405 (3 uM) (n = 3 independent experiments, >550 cells analyzed per LRRK2 genotype in each experiment). Error bars: mean +/− SEM. **(C)** Scatter plot showing low correlation between *in vitro* p-Rab8A levels and cellular p-Rab10 levels in untreated conditions. Data points for both axes were estimated from Figures 1 and 4 of Kalogeropulou et al^8^.

**Table S1:** Reported effects of LRRK2 mutations on *in vitro* kinase activity, GTPase activity, GTP binding, and pS935 level.

| **Mutation** | **Domain** | **Category**<sup>5,8,58,59</sup> | ***in vitro* kinase activity**<sup>7,8,60,61</sup> | **GTPase activity**<sup>10,44,45,59,62</sup> | **GTP binding**<sup>10,16,59,63,64</sup> | **pS935 level**<sup>7,8,16</sup> |
| --- | --- | --- | --- | --- | --- | --- |
| T1348N | ROC | Hypothesis-testing | n/a | Decrease | Decrease (GTP-binding deficient) | Decrease |
| D2017A | Kinase | Hypothesis-testing | Decreased (kinase dead) | n/a | No change | No change or decrease |
| R1398H | ROC | Protective | n/a | Increase | Decrease | No change |
| N1437H | ROC | Causal | Small decrease | Decrease | Increase | Decrease |
| R1441C | ROC | Causal | No change | Decrease | Increase | Decrease |
| R1441G | ROC | Causal | No change or small decrease | Decrease | Increase | Decrease |
| R1441H | ROC | Causal | No change | Decrease | Increase | Decrease |
| Y1699C | COR | Causal | No change or increase | Decrease | Increase | Decrease |
| R1728H | COR | Risk factor | Increase | n/a | No change | No change or small decrease |
| I2012T | Kinase | Risk factor | Decrease | n/a | No change | No change or small decrease |
| G2019S | Kinase | Causal | Increase | No change | No change | No change |
| I2020T | Kinase | Causal | Decrease, no change, or increase | No change | No change or increase | Decrease |
| T2031S | Kinase | Risk factor | Increase | n/a | No change | No change |
| G2385R | WD40 | Risk factor | Decrease, no change, or increase | n/a | No change | Decrease |

Mutant effects on *in vitro* kinase activity, GTPase activity, GTP binding, and pS935 level are reported relative to WT LRRK2. n/a: not available.

## Acknowledgements

We are grateful to all members of the Altschuler-Wu lab for their feedback. We thank Dario Alessi for kindly providing LRRK2 expression constructs, the MRC PPU Reagents and Services facility for PPM1H constructs, and Harald Stenmark for the gift of the pmCherry-2xFYVE construct. We also thank Mark von Zastrow, Kevan Shokat, and Sandra Schmid for helpful feedback throughout the course of this study. We gratefully thank support from the National Science Foundation Graduate Research Fellowship (C.R. grant number 1650113, C.S.W. grant number 2034836), a gift from an anonymous donor (C.R. and C.S.W.), the Alexander and Eva Nemeth Foundation (R.J.N.), the NCI-NIH RO1 CA184984 (L.F.W.), and CZI (L.F.W. and S.J.A.).

## Author Contributions

Conceptualization: C.R., M.P.J., L.F.W., and S.J.A.; Investigation & Visualization: C.R.; Data Analysis: C.R., C.S.W.; Microscopy Screen Data Exploration: C.R., L.R., K.K.; Resources: R.J.N.; Writing – Original Draft: C.R.; Writing – Review & Editing: C.R., M.P.J., L.F.W., S.J.A., with input from all authors; Supervision & Funding Acquisition: L.F.W., S.J.A.

## Supplementary Video Legends

**Video S1: Live imaging of R1441G LRRK2 localization upon VPS34 inhibition**. T-REx HEK293 cells expressing GFP-tagged R1441G LRRK2 (gray) were stained with Hoechst (blue). Video shows live imaging of R1441G LRRK2 after addition of SAR405 (3 uM). Images were acquired approximately every 2 min. over the course of one hour (video speed is one image per second).

**Video S2: Live imaging of G2019S LRRK2 localization upon VPS34 inhibition**. T-REx HEK293 cells expressing GFP-tagged G2019S LRRK2 (gray) were stained with Hoechst (blue). Video shows live imaging of G2019S LRRK2 after addition of SAR405 (3 uM). Images were acquired approximately every 2 min. over the course of one hour (video speed is one image per second).

**Video S3: Live imaging of WT LRRK2 localization upon VPS34 inhibition**. T-REx HEK293 cells expressing GFP-tagged WT LRRK2 (gray) were stained with Hoechst (blue). Video shows live imaging of WT LRRK2 after addition of SAR405 (3 uM). Images were acquired approximately every 2 min. over the course of one hour (video speed is one image per second).

**Video S4: Live imaging of empty vector (GFP only) localization upon VPS34 inhibition**. T-REx HEK293 cells expressing empty vector (GFP only) control (gray) were stained with Hoechst (blue). Video shows live imaging of GFP after addition of SAR405 (3 uM). Images were acquired approximately every 2 min. over the course of one hour (video speed is one image per second).

## Materials and Methods

### Antibodies and reagents

The following primary antibodies were used at the indicated dilutions for immunofluorescence (IF) and Western blot (WB): rabbit anti-Rab10 (abcam, #ab237703, 1:500 for IF, 1:1000 for WB), rabbit anti-Rab10 (phospho-T73) (abcam, #ab241060, 1:100 for IF, 1:500 for WB), rabbit anti-Rab8A (Cell Signaling Technology, #6975, 1:100 for IF, 1:1000 for WB), rabbit anti-Rab8A (phospho-T72) (abcam, ab230260, 1:1000 for WB), mouse anti-a-tubulin (Cell Signaling Technology, #3873, 1:2000 for IF), rabbit anti-a-tubulin (Cell Signaling Technology, #2125, 1:50 for IF), mouse anti-HA-tag (ThermoFisher, #26183, 1:500 for IF), mouse anti-Rab5A (Cell Signaling Technology, #46449, 1:200 for IF), rabbit anti-EEA1 (Cell Signaling Technology, #3288, 1:100 for IF), rabbit anti-Rab7 (Cell Signaling Technology, #9367, 1:100 for IF), rabbit anti-Rab11 (Cell Signaling Technology, #5589, 1:100 for IF), rabbit anti-GM130 (Cell Signaling Technology, #12480, 1:3200 for IF), rabbit anti-LC3A/B (Cell Signaling Technology, #12741, 1:200 for IF), rabbit anti-LAMP1 (Cell Signaling Technology, #9091, 1:300 for IF), mouse anti-LRRK2 (NeuroMab/AntibodiesInc, #75-253, 1:1000 for WB), mouse anti-GAPDH (Cell Signaling Technology, #97166, 1:1000 for WB). AlexaFluor secondary antibodies (ThermoFisher) were used in immunofluorescence assays, and IRDye secondary antibodies (LI-COR) were used in Western blots. We note from the abcam product page that the anti-Rab8A (phospho-T72) antibody cross-reacts with phosphorylated Rab3A, Rab10, Rab35 and Rab43.

Doxycycline (#D9891) and Blasticidin (#203350) were from Millipore Sigma. Hygromycin B (#10687010) and Zeocin (#R25001) were from ThermoFisher. Collagen type I from rat tail was purchased from Corning (#354236).

SAR405 (#S7682) and Chloroquine (#S4157) were from Selleck. MLi-2 (#19305), VPS34-IN1 (#17392), Nocodazole (#13857), Brefeldin A (#11861), Rotenone (#13995), and Docetaxel (#11637) were from Cayman Chemical. Compounds were used at the indicated concentrations.

With the exception of the initial microscopy-based survey of LRRK2 localization, DMSO was used at a final concentration of 0.1% and SAR405 was used at 3 uM.

We note that MLi-2 leads to Rab10 dephosphorylation within 10 minutes^65^. MLi-2 has been reported to cause LRRK2 dephosphorylation and enhanced microtubule association for some LRRK2 genotypes at later timepoints^8,65,66^.

### Plasmids and site-directed mutagenesis

cDNA constructs pcDNA5FRT/TO-GFP-LRRK2 WT (DU113363), pcDNA5FrtTO-GFP-LRRK2 T1348N (DU34766), pcDNA5FrtTO-GFP-LRRK2 R1441G (DU13388), and pcDNA5FrtTO-GFP-LRRK2 D2017A (DU13364) for expression of GFP-tagged, full-length LRRK2 were from Dr. Dario Alessi. Site-directed mutagenesis (SDM) was used to generate additional LRRK2 mutant plasmids. Some LRRK2 plasmids contained SNP S1647T, which was reverted to the consensus WT amino acid (S1647) via SDM prior to performing all experiments in this study with the exception of the initial microscopy screen (Fig. S1A).

SDM was performed using the Q5 Site-Directed Mutagenesis Kit (NEB #E0554S) according to the manufacturer’s directions. The NEBaseChanger web-based tool was used to generate appropriate primer sequences and annealing temperatures. For all plasmids generated via SDM, the entire LRRK2 insert was sequence-verified before use in experiments.

Additional plasmids utilized in this study: pcDNA5FRT/TO GFP (DU13156) was from Dr. Dario Alessi, pCMV5D-HA-PPM1H (DU62789) and pCMV5D-HA-PPM1H H153D (DU62928) were from the MRC PPU Reagents and Services facility, and pmCherry-2xFYVE was a gift from Harald Stenmark (Addgene plasmid # 140050). Plasmids were verified by Sanger sequencing before use in experiments. Plasmids with DU numbers can be found on the University of Dundee MRC PPU website: https://mrcppureagents.dundee.ac.uk/.

DNA constructs were amplified in high efficiency NEB 5-alpha Competent *E. coli* (#C2987H) and purified using a QIAprep Spin Miniprep Kit (#27106) or QIAGEN Plasmid Plus Midi Kit (#12943).

### Flp-in T-REx HEK293 cell line generation and cell culture

The Flp-in T-REx HEK293 system (Invitrogen) was utilized for both stable plasmid transfection as well as doxycycline-induced expression, as described previously^7,22^. Base T-REx media was DMEM supplemented with 10% FBS, 1% penicillin/streptomycin solution. Prior to transfection, the T-REx HEK293 host cell line was cultured in base T-REx media supplemented with 100 ug/mL Zeocin and 15 ug/mL Blasticidin. Upon stable integration of pcDNA5FRT/TO plasmids, T-REx HEK293 cell lines were maintained in base T-REx media supplemented with 50 ug/mL Hygromycin B and 15 ug/mL Blasticidin. Cell lines were tested for mycoplasma contamination.

Transfections of GFP-tagged LRRK2 or GFP-only control pcDNA5FRT/TO plasmids were performed using jetPRIME transfection reagent (Polyplus #101000027) in base T-REx media. Cells were co-transfected in 6-well culture plates at 60-80% confluency at a 9:1 ratio of pOG44 plasmid to pcDNA5FRT/TO plasmid of interest. Media was replaced four hours after transfection. 24 hours after transfection, cells were split into 10 cm dishes in base T-REx media. 48 hours after transfection, media was replaced with selection media (base T-REx media supplemented with 10 ug/mL Blasticidin and 100 ug/mL Hygromycin B). Transfected cells were selected for approximately two weeks to isolate cells that had successful plasmid integration into the FRT site. For experiments, expression of the indicated protein was induced using doxycycline (1 ug/mL).

### Immunofluorescence microscopy

T-REx HEK293 cells were seeded in 384-well PhenoPlates (PerkinElmer #6057302) manually coated with collagen I in T-REx base media supplemented with doxycycline (1 ug/mL). Plated cells were incubated for 48 hours before indicated drug treatment and immunofluorescence.

Cells were fixed with 4% paraformaldehyde for 15 minutes followed by three PBS washes at room temperature. Cells were then permeabilized for 10 minutes using 0.3% Triton X-100 in PBS and washed three times with PBS at room temperature. For conditions utilizing anti-LAMP1 or anti-LC3A/B antibodies, cells were alternatively fixed with ice-cold 100% methanol for 10 minutes at −20°C and not subjected to a permeabilization step. Cells were then blocked with 1% BSA in PBS-T (0.1% Tween-20 in PBS) for one hour at room temperature. Cells were incubated with primary antibodies in blocking buffer for 16-18 hours at 4°C. After three PBS washes, cells were incubated with secondary antibodies in blocking buffer for one hour at room temperature followed by three PBS washes. Hoechst (Invitrogen #H3570) was added at a 1:1000 dilution for 10 minutes to one of the final PBS washes for nuclei staining. An anti-alpha-tubulin (mouse or rabbit) antibody was included when possible to aid in cell segmentation during image processing.

Microscopy was performed using a PerkinElmer Operetta CLS System with confocal spinning-disk mode. Cells were imaged with a 40x water objective and standard filter sets.

### Live cell microscopy

T-REx HEK293 cells were seeded in 96-well PhenoPlates (PerkinElmer #6055302) manually coated with collagen I in T-REx base media supplemented with doxycycline (1 ug/mL). Plated cells were incubated for 48 hours before live imaging. To label nuclei, cells were incubated with Hoechst (Invitrogen #H3570) diluted 1:2000 in media for 10 minutes at 37°C before washing two times with warm media.

Microscopy was performed with a PerkinElmer Operetta CLS System, utilizing the live cell chamber to maintain an environment of 37°C and 5% CO2. Cells were imaged using confocal spinning-disk mode with a 40x water objective and standard filter sets. Prior to drug treatment, cells were imaged once to capture a pre-treatment state. Immediately following drug addition, cells were imaged at the indicated regular intervals for up to 60 minutes. For live imaging of MLi-2 treatment effects, imaging was briefly paused (<1 minute) to allow for MLi-2 addition after 30 minutes of imaging.

### Initial microscopy-based survey of LRRK2 localization

T-REx HEK293 cells expressing GFP-tagged LRRK2 plasmids were treated with the following compounds as follows:

| Compound | Target endo-lysosomal pathway | Concentration; treatment duration |
| --- | --- | --- |
| DMSO | N/A | 0.2%; 2 hours |
| Brefeldin A<sup>56,67</sup> | ER-Golgi transport | 25 uM; 2 hours |
| Docetaxel<sup>68-70</sup> | Microtubule transport | 2 uM; 2 hours |
| Nocodazole<sup>56,71,72</sup> | Microtubule transport | 10 uM; 2 hours |
| Rotenone<sup>73,74</sup> | Mitophagy, mitochondria function | 1 uM; 1.5 hours |
| Chloroquine<sup>19,75</sup> | Autophagy, lysosome function | 30 uM; 2 hours |
| SAR405<sup>31,76,77</sup> | Endocytosis | 2 uM; 2 hours |

Following compound treatment, cells were fixed with 4% paraformaldehyde and processed for microscopy analysis as described above.

### Dextran endocytosis assay

T-REx HEK293 cells were plated for live cell microscopy and stained with Hoechst as described above. Cells were then incubated with 10 kDa CF640R-labeled dextran (100 ug/mL, Biotium #80115) for 15 minutes at 37°C in the presence of DMSO or SAR405. After incubation, cells were washed five times with pre-warmed medium containing either DMSO or SAR405.

Cells were then immediately imaged via live-cell microscopy every 10 minutes for one hour in the appropriate media containing DMSO or SAR405.

### PPM1H and 2xFYVE transient transfections

T-REx HEK293 cells stably transfected with LRRK2 plasmids were seeded in 96-well PhenoPlates (PerkinElmer #6055302) manually coated with collagen I in T-REx base media. Cells were transfected the next day at 40-60% confluence using jetPRIME transfection reagent (Polyplus #101000027) at a 2:1 jetPRIME reagent to DNA ratio.

HA-PPM1H, HA-PPM1H H153D plasmids were transfected using 25 ng DNA. Media was replaced with T-REx base media supplemented with doxycycline (1 ug/mL) four hours after transfection. After 24 hours, drug treatment and immunofluorescence was performed.

The pmCherry-2xFYVE plasmid was transfected using 20 ng DNA. Media was replaced with T-REx base media supplemented with doxycycline (1 ug/mL) four hours after transfection. After 48 hours, drug treatment and live imaging was performed.

### Image analysis and data processing

Maximum intensity projections obtained from three to four z-slices per field of view were used for analysis and for representative images. Representative images were processed identically across LRRK2 genotypes within each figure panel. For Supp. Video 4, the range between pixel minimum and maximum values was widened compared to Supp. Videos 1-3 to more easily visualize the brighter fluorescence signal for empty vector (GFP only).

Image analysis methods used here have been previously described in detail^78,79^. All image analysis was done in MATLAB. In brief, background intensity was subtracted from maximum projection images to remove nonuniform background illumination using the NIH ImageJ rolling ball background subtraction algorithm. Individual cells in each image were then identified by an in-house watershed-based segmentation algorithm that utilizes nuclear and cytoplasmic markers to determine cell boundaries^79^. In each image channel for biomarkers of interest, numerous features were measured and extracted from each identified cell, including whole-cell morphology and intensity features as well as object-based morphology and intensity features^79^. The following single-cell features were utilized: total cell area, cytoplasmic area, nuclear area, area and intensity of marker objects in the cell, intensity of a marker in the cell (non-object based), and the Pearson correlation coefficient across image channels within the cytoplasm of the cell (non-object based). For markers of interest in which object-specific features were desired (LRRK2, Rab5A, and 2xFYVE), objects were identified as follows: First, foreground images were calculated for the channel of interest by applying a tophat filter and gaussian blur to reduce pixel noise^80^. An object mask was then calculated by applying a threshold value to the foreground image. The threshold value was determined for each cell by first calculating the median pixel intensity of the marker of interest in that individual cell (calculated from the original, unfiltered cell image). The final object threshold was then calculated as the median intensity raised to an exponent and multiplied by a constant. The scaling values (exponent and constant multiplier) were held constant across all cells analyzed in all experiments for the given biomarker. The scaling values were empirically selected for each biomarker to allow effective object identification across a range of object and cytoplasm intensities.

Quality control was performed before final analysis. Poorly segmented cells were removed according to the following parameters: cytoplasmic to nuclear area ratio less than 0.5 or total cell area less than 2500 px^2^ (~225 um^2^). For HA-PPM1H and HA-PPM1H H153D transient transfection experiments, unsuccessfully transfected cells had a mean cellular HA-tag fluorescence of less than 80 a.u. and were excluded from analysis.

“Percent of cells with LRRK2 on objects/endosomes” was calculated as the number of cells in which the total area of LRRK2 objects in the cell was greater than 40 px^2^ (~3.5 um^2^) divided by the total number of cells. (The minimum area threshold was determined empirically to minimize false positives due to artefacts or small cytoplasmic objects previously described for some LRRK2 mutants^7,25^. Nevertheless, a small fraction of DMSO-treated cells had cytoplasmic objects that surpassed this threshold (see Fig. 1C, Fig. S1E).) The term “endosomes” was used instead of “objects” in plot labels where appropriate after determining the identity of VPS34 inhibition-induced LRRK2^+^ endosomes.

### Cell lysate preparation and western blotting

Western blot methods were roughly based on a published protocol for LRRK2 immunoblotting^81^. T-REx HEK293 cells were plated in 6-well tissue culture plates in T-REx base media supplemented with doxycycline (1 ug/mL). After 48 hours, cells were treated with the indicated compounds for 30 minutes before cell lysis for 30 minutes on ice. Lysis buffer contained the following components: 50 mM Tris HCl pH 7.5, 1% Triton X-100, 0.27 M sucrose, 1 mM EGTA, 1 mM EDTA, 2 mM PMSF Protease Inhibitor (ThermoFisher #36978), 1 ug/mL Microcystin-LR (Cayman Chemical #10007188), and 1.5X Halt Protease and Phosphatase Inhibitor Cocktail (ThermoFisher #36978). Cell extracts were centrifuged at 16,500 x g for 15 minutes at 4°C to clarify the lysate and protein concentrations were measured via BCA assay (Pierce).

Cell extracts (15 ug) were resolved by SDS-PAGE using 4-15% TGX precast gels (Bio-Rad #4561086) with tris/glycine/SDS buffer and transferred onto PVDF membranes using the Bio-Rad Mini-PROTEAN Electrophoresis System. Membranes were blocked with LI-COR blocking buffer for one hour at room temperature followed by incubation with primary antibodies diluted in blocking buffer overnight at 4°C. Membranes were washed thoroughly with TBS-T (0.1% Tween-20 in TBS) before incubation with secondary antibodies (IRDye, LI-COR) diluted in blocking buffer at room temperature for one hour. Membranes were washed in TBS-T before imaging with an Azure 600 imager (Azure Biosystems).

### Statistical analysis and plotting

Statistical analyses and plotting were performed in R (version 4.0.3) using standard packages (ggplot2, stats, DescTools). For experiments with only two groups, one-tailed or two-tailed unpaired t-tests were performed with Bonferroni correction to adjust for multiple comparisons where appropriate. For experiments comparing three or more groups, a one-way ANOVA (analysis of variance) was used. If the ANOVA determined that there was a difference between the means of at least two of the treatment groups, a post hoc test was used. Dunnett’s post hoc test was used when determining statistical significance for comparisons between individual experimental groups and a control group, while Tukey’s post hoc test was used when determining statistical significance for comparisons between all individual treatment groups. Cartoons in Fig. 4E and Fig. S2B were created using BioRender.com.

